# Predictors of Malaria Rapid Diagnostic Tests’ Utilisation Among Healthcare Workers in Zamfara State

**DOI:** 10.1101/363697

**Authors:** Usman Rabi, Ahmad A. Umar, Saheed Gidado, A.A Gobir, Izuchukwu F. Obi, IkeOluwapo Ajayi, Olufemi Ajumobi

**Affiliations:** Nigeria Field Epidemiology and Laboratory Training Programme, Abuja, Nigeria; Department of Community Medicine, Ahmadu Bello University, Zaria, Nigeria; African Field Epidemiology Network, Abuja, Nigeria; Department of Epidemiology and Medical Statistics, University of Ibadan, Ibadan, Nigeria; Department of Community Medicine, University of Nigeria Teaching Hospital Enugu, Nigeria

**Keywords:** Malaria rapid diagnostic test, knowledge, utilisation, healthcare worker, Nigeria

## Abstract

**Introduction:** Early diagnosis and prompt and effective treatment is one of the pillars of malaria control Malaria case management guidelines recommend diagnostic testing before treatment using malaria Rapid Diagnostic Test (mRDT) or microscopy and this was adopted in Nigeria in 2010. However, despite the deployment of mRDT, the use of mRDTs by health workers varies by settings. This study set out to assess factors influencing utilisation of mRDT among healthcare workers in Zamfara State, Nigeria.

**Methods:** A cross-sectional study was carried out among 306 healthcare workers selected using multistage sampling from six Local Government Areas between January and February 2017. Mixed method was used for data collection. A pre-tested self-administered questionnaire was used to collect information on knowledge, use of mRDT and factors influencing utilization. An observational checklist was used to assess the availability of mRDT in the six months prior to this study. Data were analyzed using descriptive statistics such as means and proportions. Association between mRDT use and independent variables was tested using Chi square while multiple regression was used to determine predictors of use at 5% level of significance.

**Results:** Mean age of respondents was 36.0 ± 9.4years. Overall, 198 (64.7%) of health workers had good knowledge of mRDT; malaria RDT was available in 33 (61.1%) facilities. Routine use of mRDT was reported by 253 (82.7%) healthcare workers. This comprised 89 (35.2%) laboratory scientists/technicians, 89 (35.2%) community health extension workers/community health officers; 59 (23.3%) nurses and 16 (6.3%) doctors. Predictors of mRDT utilisation were good knowledge of mRDT (adjusted OR (aOR):3.3, CI: 1.6-6.7), trust in mRDT results (aOR: 4.0, CI: 1.9 - 8.2), having being trained on mRDT (aOR: 2.7, CI: 1.2 - 6.6), and provision of free mRDT (aOR: 2.3, CI: 1.0 - 5.0).

**Conclusion:** This study demonstrated that healthcare worker utilisation of mRDT was associated with health worker and health system-related factors that are potentially modifiable. There is need to sustain training of healthcare workers on benefits of using mRDT and provision of free mRDT in health facilities.

## Introduction

Malaria remains a major public health problem in many countries of the world. Despite the progressive reduction in malaria cases and deaths, it is estimated that an estimated 216 million cases of malaria occurred worldwide in 2016 with 90% from the African region.^1^ Fifteen countries accounted for 80% of the 445,000 malaria deaths worldwide; these countries are all in sub-Saharan Africa which include Nigeria^1^.

In 2016, Nigeria accounted for more than 50% of all malaria cases in sub-Saharan Africa^1^; the disease is responsible for two-thirds of outpatient visits to health facilities, one-third of childhood deaths, one-quarter of deaths in children under one year and 11% maternal deaths. The financial loss due to malaria annually is estimated to be about 132 billion naira in form of treatment costs, prevention and loss of man-hours among others; yet, it is a treatable and preventable disease^2^. Malaria prevalence in Zamfara State has remained consistently high, 69.9% ^3^ with less than one percent of children with fever being tested for malaria^4^.

The WHO in 2010, recommended confirmation of malaria in febrile illness prior to treatment with artemisinin combination therapy^5^. Attaining the objective of test and treat for all suspected malaria cases using RDT or microscopy^6^ make it imperative for all health workers to have access to, and appropriately utilize malaria diagnostic tools. Although microscopy is recognized as the gold standard in malaria diagnosis, it has been limited in availability, often of poor quality, time-consuming, labor-intensive, and costly^7,8^ especially in resource-poor settings. Lack of equipment, reagents, and expertise for malaria microscopy in the majority of peripheral health centers and the constant power supply has equally limited its use. More so, presumptive diagnosis based on malaria symptoms has proven to be unspecific^9–11^. These shortcomings of microscopy and presumptive diagnosis have favored the deployment and use of mRDTs which have been found to be cost-effective^12–14^ and allow diagnosis even in health settings lacking any laboratory facility. Malaria RDT use is expected to not only improve malaria management but also limit malaria treatment costs^15^. Deployment to mRDT to health facilities commenced in Nigeria in the year 2007.

Factors such as heavy workload, lack of trust, cost, training on the use of RDTs have been considered to influence RDT use^16–21^. A study reported a high proportion (61.5%) of healthcare workers perceived mRDTs as unreliable, one-third (30.8%) of healthcare workers had supply issues with mRDT, 15.4% of them reported a preference for other methods of malaria diagnosis and one-fifth (26%) of healthcare workers were ignorant about mRDT.^16^ These factors are generic and may vary in different settings.

There is a paucity of data concerning the mRDT use and factors influencing utilisation among healthcare workers in Zamfara State. Lack of malaria testing could impair the ability of health workers to make informed and prompt treatment decision based on parasitological diagnosis^5^. This study aimed to investigate the knowledge of mRDT, mRDT availability and use as well as factors influencing mRDT utilisation in health facilities in Zamfara State.

## Methods

### Study area

The study was conducted in Zamfara State, North West Nigeria. The State has a projected population of 4,466,775 (based on the 2006 Census population with an annual growth rate of 3.2%). The climate of Zamfara is tropical with a temperature rising up to 38 °C (100.4 °F) and above between March to May. The state experiences malaria transmission all year-round with peak transmission during the rainy season between May and September. The State operates a three-tier healthcare delivery services namely primary, secondary (General Hospitals) and tertiary spread across urban and rural areas. The State has a total of 712 health facilities distributed across 14 Local Government Areas (LGAs). These health facilities are as follows; 71 Primary Health Centres, 607 Health Clinics, 10 private hospitals, 22 General Hospitals, 1 Specialist Hospital and 1 Federal Medical Center. The State has a total of 3,458 healthcare workers working in these health facilities. Majority of the facilities in the State offer malaria diagnosis and treatment services^22^. Generally, trained staff of public primary health centers offer malaria diagnosis using mRDT while trained laboratory scientists at public general hospitals (secondary care level) offer both malaria microscopy and mRDT services. The State has benefitted from several Malaria intervention programs over the years such as Partnership for Reviving Routine Immunization in Northern Nigeria-Maternal and Neonatal Child Health (PRRINN/MNCH), Malaria Action Program for States (MAPS) and of recent, the STOP/Malaria Frontline project to improve the effectiveness of malaria control in Zamfara State. A crosssectional study was carried out among health workers. in public health facilities in the state between January and February 2017.

### Sample size and sampling technique

A sample size of 306 was calculated using sample size formula for single proportion;

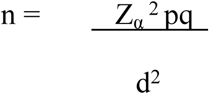

Where:
the p= proportion of health workers that use malaria RDT to diagnose malaria in public health facilities, (0.85)^23^

q= 1-p= 1-0.852= 0.148

d= level of precision, 0.05

α = level of significance, 5%

Z_α_= standard normal deviate, 1.96

A three-stage sampling technique was used to select study respondents. Two (2) LGAs were randomly selected by balloting from each of the three senatorial zones of the State giving a total of 6 LGAs namely; Kaura Namoda and Zurmi LGAs (Zamfara North zone), Gusau and Bungudu LGAs (Zamfara Central zone), Anka and Talata Mafara LGAs (Zamfara West zone). List of all public health facilities from the selected LGAs based on the level of care was stratified into primary and secondary facilities. Eight Primary Health Care centers (PHCs) were selected from each of the selected LGA by balloting giving a total of 48 PHCs while the General hospital in each of the LGA selected was purposively selected for the study. However, where there was more than one General Hospital in a selected LGA, one was selected by balloting. This gave an overall total of 54 health facilities selected for the study. A sampling frame of all healthcare workers was developed using the facility’s nominal roll. Health workers were selected by stratified sampling proportionate to size until required sample size was obtained.

### Data collection

Six trained research assistants distributed the questionnaires. Semi-structured self-administered questionnaires were used to obtain information on respondents’ socio-demographic characteristics, knowledge of mRDT, malaria diagnostic methods used in health facilities, utilization of malaria RDT among health care workers, training on malaria case management, supervision on malaria RDT use and factors affecting malaria RDT utilization. The research assistants administered health facility observational checklists to assess the availability of mRDTs at the facilities within the last six months.

### Data processing and analysis

Questionnaires were manually checked for completeness and consistency with corrections made daily. Data were entered, cleaned and analyzed using Epi-info Version 7. Data were summarized using descriptive statistics such as means and standard deviations for quantitative variables such as age, years of practice and knowledge score while frequencies and proportions were generated for categorical variables (the cadre of health worker, the proportion of febrile patients who get tested using mRDT, mRDT availability, and mRDT use). Results of the analysis were presented in tables and charts. Healthcare workers’ mRDT knowledge scores were calculated thus; 8 questions evaluated knowledge of mRDT, each correct answer was given a score of 1 and an incorrect answer was given a score of 0. Total scores were computed for each respondent and converted into percentages. A score of less than 50% was graded as poor knowledge, between 50% and 75% as fair knowledge and greater than 75% as good knowledge. Bivariate analysis was used to test the association between categorical dependent and independent variables. Those significant at p-value ≤ 5% were put in the logistic regression model to control for confounders to determine predictors of mRDTs by healthcare workers. Odds Ratios and 95% Confidence Intervals (CIs) were presented.

### Ethical considerations

This research was granted ethical approval by the Ethics and Research Committee of Zamfara State Ministry of Health (Reference number-ZSHREC/03/10/2016). Participation was voluntary and written informed consent was obtained from all respondents. The participants were not at any point in time exposed to harm and were free to opt out at any time during the interview. Confidentiality of collected information was maintained by using unique non-personal identifier codes for the respondents. The completed questionnaire was kept under lock and key.

## Results

### Characteristics of respondents

Overall, 306 healthcare workers participated in the study and their mean age was 36.0yrs, SD: 9.4yrs. Most, 128 (41.8%), of the respondents were aged 25 to 34years. They were mostly males, 204 (66.7%). CHEWs represented 105 (34.3%) of respondents and 21 (6.9%) were doctors. Most were married (78.1%, n = 239). The average duration of practice was 11.0 ± 9.1yrs (Table 1).

**Table 1:** Frequency ddistribution of socio-demographic characteristics of the respondents (N = 306)

| **Characteristics** | **Frequency (%)** |
| --- | --- |
| **Age group (in years)** |  |
| <25 | 22 (7.2) |
| 25-34 | 128 (41.8) |
| 35-44 | 90 (29.4) |
| 45-54 | 54 (17.7) |
| ≥55 | 12 (3.9) |
| **Sex** |  |
| Male | 204 (66.7) |
| **The cadre of healthcare worker** |  |
| Doctor | 21 (6.9) |
| Nurse/Midwife | 83 (27.1) |
| Laboratory Scientist/Technician | 97 (31.0) |
| CHEW/CHO | 105 (34.3) |
| **Marital Status** |  |
| Single | 63 (20.6) |
| Married | 239 (78.1) |
| Widowed | 4 (1.3) |
| **Duration of practice (years)** |  |
| 1-5 | 95 (31.1) |
| 6-10 | 95 (31.1) |
| 11-15 | 47 (15.4) |
| 16-20 | 23 (7.5) |
| 21-25 | 8 (2.6) |
| >25 | 38 (12.4) |

### Health workers’ knowledge of mRDT

Among the 306 health workers, 245 (80.1%) knew the meaning of mRDT, while 236 (77.0 %) knew what mRDT assesses. All the respondents, 306 (100%) knew that blood was used for the test. Two hundred and eighty-two (92.0%) knew how to carry out mRDT. Mean knowledge score for respondents was 82.0 (standard deviation (SD): 15.8). Overall, more than half of respondents (64.7%, n = 198) had good knowledge of mRDT. According to cadre, 71.1% of laboratory scientists/technicians, 63.8% of CHEWs/CHOs, 61.9% of doctors and 59.0% nurses had good knowledge of malaria RDT. (Table 2).

**Table 2:** Level of knowledge of malaria Rapid Diagnostic Test by Professional Cadre of Respondents in Selected Health Facilities, Zamfara State.

| Professional cadre | Knowledge grade |  |  |  |  |  | Total<br>N = 306 |
| --- | --- | --- | --- | --- | --- | --- | --- |
|  | Good<br>N = 198 |  | Fair<br>N = 99 |  | Poor<br>N = 9 |  |  |
|  | Frequency | % | Frequency | % | Frequency | % |  |
| Doctors | 13 | 61.9 | 4 | 19.1 | 4 | 19.1 | 21 |
| Nurse/Midwife | 49 | 59.0 | 32 | 38.6 | 2 | 2.4 | 83 |
| CHEW/CHO | 67 | 63.8 | 36 | 34.3 | 2 | 1.9 | 105 |
| Lab. Scientist/Technician | 69 | 71.1 | 27 | 27.8 | 1 | 1.0 | 97 |

### Availability and use of mRDT among healthcare workers

Thirty-three (61.1%) out of the 54 of health facilities had mRDT in stock. Overall, 253 (82.7%) of the healthcare workers reported using malaria RDT routinely before making a diagnosis of malaria. This comprised 89 (35.2%) laboratory scientists/technicians, 89 (35.2%) community health extension workers/community health officers; 59 (23.3%) nurses and 16 (6.3%) doctors. Nurses/Midwives (OR: 2.7, 95% CI: 1.5 – 5.0) and Laboratory scientists/technicians (OR: 3.1, 95% CI: 1.4 – 6.8) were significantly more likely to use mRDT compared to doctors and CHEWs/CHOs (Table 3).

**Table 3:** Utilisation OF Malaria Rapid Diagnostic Test Among Healthcare Workers in Selected Health Facilities, Zamfara State (N=253)

| **Professional cadre** | **n (%)** | **OR (95%CI)** | **p-value** |
| --- | --- | --- | --- |
| Doctor | 16 (6.3) | 1.5 (0.5 - 4.4) | 0.606 |
| Nurse/Midwife | 59 (23.3) | 2.7 (1.5 - 5.0) | 0.002 |
| CHEW/CHO | 89 (35.2) | 0.8 (0.4 - 1.5) | 0.592 |
| Lab. Scientist/Technician | 89 (35.2) | 3.1 (1.4 - 6.8) | 0.007 |

### Factors influencing mRDT use

Healthcare workers with good knowledge of mRDT were 2.7 times more likely to use it (p = 0.002). Those who have had training in malaria case management had 2.4 times odds of using mRDT (p = 0.003). Healthcare workers in facilities that do mRDT for free were 2.8 times more likely to use it (p = 0.036) compared to facilities where patients have to pay for it. Healthcare workers who have trust in mRDT, have had training on mRDT and those who have received supportive supervision ((p = 0.002) were 3 times more likely to use mRDT compared to those who did not have training and did not receive supportive supervision. Predictors of mRDT utilisation include health workers’ having good knowledge of mRDT (aOR: 3.3, 95% CI: 1.6 – 6.7), trust in mRDT results, (aOR: 4.0, 95% CI: 2.0 – 8.3), training on mRDT (aOR: 2.8, 95% CI: 1.2 – 6.7) and provision of free mRDT (aOR: 2.3, 95% CI: 1.0 – 5.0), Table 4.

**Table 4:** Association between respondents’ factors, health system factors, and utilization of malaria Rapid Diagnostic Test, Zamfara.

| Characteristic | Crude OR | p-value | aOR | p-value |
| --- | --- | --- | --- | --- |
| **Age (in years)** |  |  |  |  |
| <36 | 1.5 (0.5 – 1.7) | 0.9798 |  |  |
| >36 |  |  |  |  |
| **Sex** |  |  |  |  |
| Male | 2.5 (1.4 – 4.5) | 0.0046 |  |  |
| **Professional cadre** |  |  |  |  |
| Doctor | 1.5 (0.5 – 4.4) | 0.6062 |  |  |
| Nurse/Midwife | 2.7 (1.5 – 5.0) | 0.0019 |  |  |
| CHEW/CHO | 0.8 (0.4 – 1.5) | 0.5916 |  |  |
| Lab. Scientist/Technician | 3.1 (1.4 – 6.8) | 0.0070 |  |  |
| **Duration of practice (in years)** |  |  |  |  |
| <15 | 1.0 (0.5 – 1.8) | 0.9496 |  |  |
| ≥15 |  |  |  |  |
| **Knowledge of mRDT** |  |  |  |  |
| Good knowledge | 2.7 (1.5 – 4.9) | 0.0020 | 3.3 (1.7 – 6.7) | <0.001 |
| Poor knowledge |  |  |  |  |
| **Training on Malaria case management** |  |  |  |  |
| Trained | 2.4 (1.7 – 6.0) | 0.0003 |  |  |
| Not trained |  |  |  |  |
| **Trust in mRDT result** |  |  |  |  |
| Trust results | 3.2 (1.7 – 6.0) | 0.0070 | 4.0 (2.0 – 8.3) | <0.001 |
| Do not trust results |  |  |  |  |
| **Had training on mRDT in the last 6 months** |  |  |  |  |
| Trained | 3.4 (1.7 – 6.6) | 0.0003 | 2.8 (1.2 – 6.7) | 0.040 |
| Not trained |  |  |  |  |
| **Received supportive supervision in the last 6 months** |  |  |  |  |
| Supervised | 3.2 (1.8 – 5.9) | 0.0002 |  |  |
| Not supervised |  |  |  |  |
| **mRDT availability** |  |  |  |  |
| Available | 1.7 (0.6 – 4.8) | 0.7917 |  |  |
| Not available |  |  |  |  |
| **Cost of mRDT** |  |  |  |  |
| Free | 2.8 (1.2 – 6.6) | 0.0363 | 2.3 (1.0 – 5.0) | 0.040 |
| Not free |  |  |  |  |

## Discussion

The study showed that healthcare workers have good knowledge of mRDT similar to a study conducted in Southeast Nigeria where 61.1% of respondents knew about mRDT.^24^ The proportion of healthcare workers who knew the meaning of mRDT was found to be higher than that in a study carried out in the six geo-political zones of Nigeria where 70% reported knowing the meaning of mRDT^21^. This is probably because of investment by government and non-governmental organizations in awareness creation on parasitological testing through training on mRDT^21^ in Zamfara state.

Majority of healthcare workers used malaria RDT routinely before making a diagnosis of malaria. This finding is similar to a study in Ogun State^25^ and systematic review of mRDT use in sub-Saharan Africa that reported a high percentage of healthcare workers used mRDT prior to administration of ACTs. However, a previous study found that doctors and laboratory technicians more likely to use mRDT compared to nurses and CHEWs/CHOs^24^. High use of mRDT among laboratory scientists is not surprising as their primary responsibility is to carry out tests.

It is widely established that the key factor in improving diagnosis of malaria is the availability of mRDTs in health facilities.^26^ This study found that rapid diagnostic test kits were available in more than half of health facilities, higher than what was reported in a study in Enugu State where 31% of health facilities had mRDT^17^ and another study in Ogun State that reported mRDT was available in 50.7% of health facilities^25^. This, however, is less than the WHO average availability target in public and private health facilities as availability is said to be inadequate if it falls below 80%. This finding underscores the need to scale-up mRDTs availability in health facilities in the State since currently, mRDTs are supplied free of charge by the government to only public health facilities. Widespread provision of malaria RDTs will play a significant role in reducing the persistent problem of malaria over-diagnosis and contribute to reduced risk of malaria under-treatment. Factors that were found to influence mRDT use in this study are similar to those found in previous studies that have reported that trust, training, and cost of mRDT affect its use^16–21^. Positive influence of healthcare workers’ trust in mRDT use in this study differs from a previous study that reported low use of mRDT despite availability because they do not trust the results^24^. This is probably because the study was conducted during the early stage of introducing mRDT into the country compared to the present day where awareness and training on mRDT have improved.

Another factor influencing mRDT utilization found in this study was training. This is similar to previous studies that showed that training of healthcare workers on mRDT improves healthcare workers’ performance with an increased likelihood of adherence to malaria treatment guidelines.^28–30^ This study also found that healthcare workers are more likely to use mRDT if the cost is free. This is similar to a previous study that reported a large improvement in the proportion of patients appropriately treated at a low cost with the introduction of mRDTs^5^. A possible reason for this is the fact that patients won’t incur any cost if they are asked to do mRDT since it is free. This, in turn, will encourage its use in health facilities, thereby, increasing the proportion of patients with the parasitological diagnosis.

### Limitations

The questionnaire captured self-reported information, hence relied primarily on respondents providing the right information. There might have been some reporting bias with probably the tendency to overestimate utilization of mRDTs in this study since this is a desirable outcome. However, this was minimized by ensuring that participants were assured of a high degree of confidentiality.

## Conclusion

The high proportion of health workers with good knowledge of mRDT in Zamfara state is commendable and could be reflection of the training that has been held in the state by multiple agents in the past. This also influenced the high use of the diagnostic kit. The drivers of mRDT use in this study (knowledge, trust, training, and provision of free mRDTs) are plausible and a good index to inform intensified efforts at capacity building of healthcare workers. The government and collaborating partners with interest in malaria control should, therefore, sustain the training of healthcare workers on mRDT and supply of free mRDTs in the health facilities in Zamfara state and the country as a whole.

## Acknowledgements

We thank the participating medical staff of all health institutions visited and the research assistants. We are grateful to the Nigeria Field Epidemiology and Laboratory Training Programme (NFELTP) for the support provided for this research. This study was supported by Cooperative Agreement Number U2GGH001876 funded by the United States Centers for Disease Control and Prevention through African Field Epidemiology Network to NFELTP. Its contents are solely the responsibility of the authors and do not necessarily represent the official views of the United States Centers for Disease Control and Prevention or the Department of Health and Human Services.

## Author’s contributions

RU and SG conceptualised the study. AAU and OA contributed to design of the study and the data tools. RU designed the study, performed the field work, data analysis and interpretation and wrote the draft manuscript. AAU, AAG, IFO, IA and OA contributed to data interpretation and provided technical inputs during the manuscript writing.

